# Effects of working memory load and CS-US intervals on delay eyeblink conditioning

**DOI:** 10.1101/2020.12.13.422606

**Authors:** Leila Etemadi, Dan-Anders Jirenhed, Anders Rasmussen

**Affiliations:** Neural Basis of Sensorimotor Control, Department of Experimental Medical Science, Lund, Sweden; Associative Learning, Department of Experimental Medical Science, Lund, Sweden; Erasmus Medical Center, Department of Neuroscience, Rotterdam, Netherlands

## Abstract

**Background:** Eyeblink conditioning is used in many different species to study motor learning and make inferences about cerebellar function. However, considerable discrepancies in performance between different species combined with evidence that awareness of stimulus contingencies affects performance indicates that eyeblink conditioning in part reflects activity in non-cerebellar regions. This questions whether eyeblink conditioning can be used as a pure measure of cerebellar function in humans.

**Methods:** Here we explored two ways to reduce non-cerebellar influences on performance in eyeblink conditioning: (1) using a short interstimulus interval, and (2) having participants do working memory tasks during the conditioning. Data were analyzed, and the influence of the interstimulus interval and working memory tasks was assessed using a linear mixed effects model.

**Results:** Our results show that subjects trained with a short interstimulus interval (150ms and 250ms) produce few conditioned responses after 100 trials. For subjects trained with a longer interstimulus interval (500ms), those who did working memory tasks produced fewer conditioned responses and had a more gradual learning curve – more akin to those reported in the animal literature.

**Conclusions:** Our results suggest that having subjects perform working memory tasks during eyeblink conditioning can be a viable strategy to reduce non-cerebellar interference in the learning.

## Introduction

Classical conditioning of the eyeblink response is a widely used experimental paradigm for the study of associative learning. In eyeblink conditioning, a subject learns to blink in response to a conditional stimulus (CS), such as a tone that is repeatedly followed by an unconditional corneal stimulus (US). The timing of the conditional response is determined by the CS–US interval used during training, such that the closure of the eyelid occurs just prior to the expected US delivery, thus protecting the eye. A short CS-US interval produces a learned blink response with a short latency, while a long CS-US interval will induce learning of a blink response with a longer latency.

In mammalian species, e.g. rabbits, cats, ferrets, rats, and mice, the associative memory trace formed during delay eyeblink conditioning, is located in the cerebellum [1–6]. Indeed, animals can acquire conditioned responses even if their entire cerebrum has been lesioned [7]. This, however, does not mean that the cerebrum cannot play a role in eyeblink conditioning. Indeed, evidence indicates that extra-cerebellar brain regions are active during eyeblink conditioning. Trace conditioning for example, where there is a temporal gap between the end of the CS and onset of the US, seems to be dependent on the hippocampus and/or cerebrum [8]. Likewise, neuroimaging data from humans undergoing eyeblink conditioning shows activity, not just in the cerebellum, but also in the cerebrum [9].

In order to relate cerebellar neurophysiological mechanisms to eyeblink conditioning in both animal and human subjects, it is desirable to reduce cerebral influence as much as possible. In this context, eyeblink responses generated by the cerebellum during and after conditioning, could be considered *true* CRs, while eyeblinks reliant on cerebral processes, e.g. voluntary blinks, may be considered false positive blinks.

Greater cortical involvement in humans could explain the glaring discrepancy between learning curves seen in the animal literature versus reports on human subjects. In the human literature, it is not uncommon to see high rates of responding already in first few blocks of trials. This is often followed by only modest increases in responding as training progresses [10–12]. Learning curves in animals, by contrast, exhibit a more gradual increase in the CR probability, and it is rare to see significant learning in the first few blocks [3,13,14]. Here we tested two strategies to reduce cerebral influence on conditioning in human subjects. The first strategy was to use a short CS-US interval in order to minimize the time for subjects to respond voluntarily. The second strategy was to reduce cortical contributions to the learning by presenting concurrent working memory tasks during conditioning. We chose working memory tasks that previous research suggest occupies the cortex so that any learned blinking to the CS would likely be cerebellum dependent only.

## Methods and Materials

### Ethical approval

In the beginning of each experiment, subjects signed a written consent form stating that they had been informed about the procedure in general terms, i.e., that their blink responses would be recorded and that they would be presented with tones and air puffs aimed at the eye. As a token of gratitude, all participants received a cinema ticket at the end of the experiment, regardless of whether they had completed the full protocol or not. The study was approved by the local ethical committee in Lund, Sweden (2017-785).

### Subjects

Subjects were 42 students (females=22; males=20) at Lund University. The age range was 24.6 ± 5.38 years (mean ± SD). The participants were divided into four groups (see Table 1). Three groups were trained with three different CS-US intervals: 150, 250 and 500 ms. The fourth group was trained with a 500 ms interval, but in addition to conditioning, subjects were instructed to concurrently perform working memory (WM) tasks.

**Table 1.**
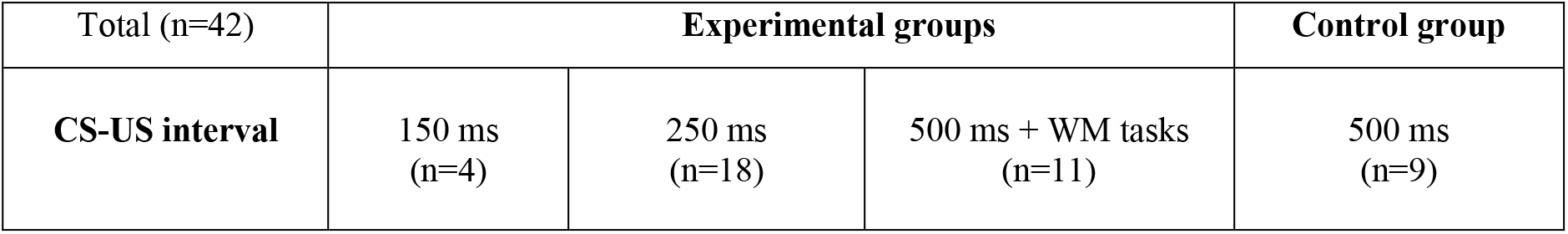
Subjects (total=42) in each CS-US interval group, and the group performing working memory (WM) tasks concurrent.

| Total (n=42) | Experimental groups |  |  | Control group |
| --- | --- | --- | --- | --- |
| CS-US interval | 150 ms<br>(n=4) | 250 ms<br>(n=18) | 500 ms + WM tasks<br>(n=11) | 500 ms<br>(n=9) |

### Eyeblink conditioning

The experiments were conducted in a quiet room at campus. First, we attached a magnet on the subject’s right eyelid and then we adjusted the intensity of the air puff so that it reliably elicited a reflexive blink response but did not cause irritation of the eye. The auditory stimulus, a 1 kHz tone of 1 s duration, was presented through a loud speaker placed approximately 1 m from the subject. The volume of the tone was adjusted to be clearly audible but not unpleasant.

Each individual received a total number of 100 trails (10 blocks of 10 trials). Of these 100 trials, 25% were probe trials meaning that the CS was presented alone. For further details on the procedure and conditioning protocol, see Rasmussen and Jirenhed [15]. In the groups that did not perform WM tasks, subjects were asked to choose a favorite TV show to watch online on a laptop during the conditioning session. The subject was asked to concentrate on the program, rather than on the presentation of tones and air puffs. In the group that received WM tasks during conditioning, subjects were told to focus fully on the working memory tasks and that the purpose of the experiment was to see the effects on WM performance of distracting stimuli in the form of air puffs and. The goal was to have the subject perceive the experiment to be a test of working memory and to be unaware that it was in fact an eyeblink conditioning experiment.

### WM tasks

A sub-set of the participants were given demanding WM tasks to perform during the eyeblink conditioning session. We selected some of the most common WM tests in use: Corsi, mental rotation, multitask, n-back (3-back), Navon, Stroop, visual search, and the Wisconsin Card Sorting task. The WM tests were run on PsyToolkit online software (PsyToolkit is developed by and belongs to Professor Gijsbert Stoet and is available at www.psytoolkit.org). Each WM task started with on-screen instructions provided by PsyToolkit. Complementary information was presented verbally when requested by subjects. Each test started with a few training examples (provided by the software).

### Materials

To detect eyelid movements, a small round neodymium magnet (diameter: 3 mm; thickness: 1 mm) was attached on the subject’s left eyelid using double stick tape. The resulting changes in the magnetic field were recorded using a sensitive GMR chip (AAH002-02E, NVE Corporation). The GMR chip and the nozzle delivering the air puff were attached to the right side of a pair of glasses that the subject wore during the test. The GMR sensor data was sampled at 1000Hz and the data was transferred to the computer via a Micro1401 AD converter (Cambridge Electronic Design). The Micro 1401 was also used to trigger the loudspeakers playing the tones and the opening of the D132202 solenoid valve (Aircom) releasing the air puff. At the beginning of the test we verified that the subject could hear the tone and that it did not produce reflex responses like blinking or flinching. The pressure of the air puff was adjusted to ensure that it produced a clear blink response without causing irritation (range from 0.5 to 1 bar).

### Data analysis

Eye-blink data was collected using the Spike2 software (CED). The data stored from WM task performances was saved in Microsoft Excel format. All data was subsequently exported to and analyzed in Matlab (Mathworks). Using custom Matlab scripts, we categorized each trial as (1) CR; (2) no CR; or (3) invalid trial. If a CR was present, the script estimated the onset and the peak of the response. All sweeps were checked manually to ensure that the script had made correct categorizations. Any errors were corrected manually. For the analysis of WM test performance, we chose two different variables: reaction time (RT) and success in the test (correct answers in %) per task/ per person. The rank was computed, using the rank function from Microsoft Excel software and an average rank presented one value for each variable of RT and success in test per person.

## Results

To test the effects of training, the interstimulus interval (ISI), and working memory tasks on the percentage of conditioned responses, we modelled CR percentage using a linear mixed effects model. A linear mixed effects model was used rather than a repeated measures analysis of variance because it is statistically more robust, takes into account individual differences, and because it copes with missing data points [16,17]. As fixed effects we used the training block (1-10), ISI (150, 250, or 500ms), sex (male or female), and whether or not the subject did working memory tasks (yes or no). We also included subject ID as a random effect in the model. The model was built in MATLAB using the fitlme function with the formula: CRs ~ 1 + Block + Sex + WM + ISI + (1 | Subject).

The model shows that block, ISI, and WM tasks has significant effects on the percentage of conditioned responses. For each block of 10 trials the CR percentage increases by an average of 2.15%. Over 10 blocks, this translates to 2.15*10 = 21.5%. This change is statistically significant (t = 8.4339, p = 4.4686e-16***, CI = 1.65 – 2.65%). Similarly, as illustrated in Figure 1A, the interstimulus interval also has a considerable effect on the CR percentage. According to our linear mixed effects model a 1ms increase in the ISI results in a 0.17% increase in the percentage of CRs. When switching from a 150ms ISI to a 500ms ISI this translates to a 0.17*350 = 59.5% increase in CR percentage. This effect is also significant (t = 10.30, p = 1.74e-22***, CI = 0.13 – 0.20%). However, contrary to our previous findings [10], sex did not have a significant effect on CRs (t = 1.621, p = 0.11, CI = −1.67 – 17.4%).

**Figure 1.**
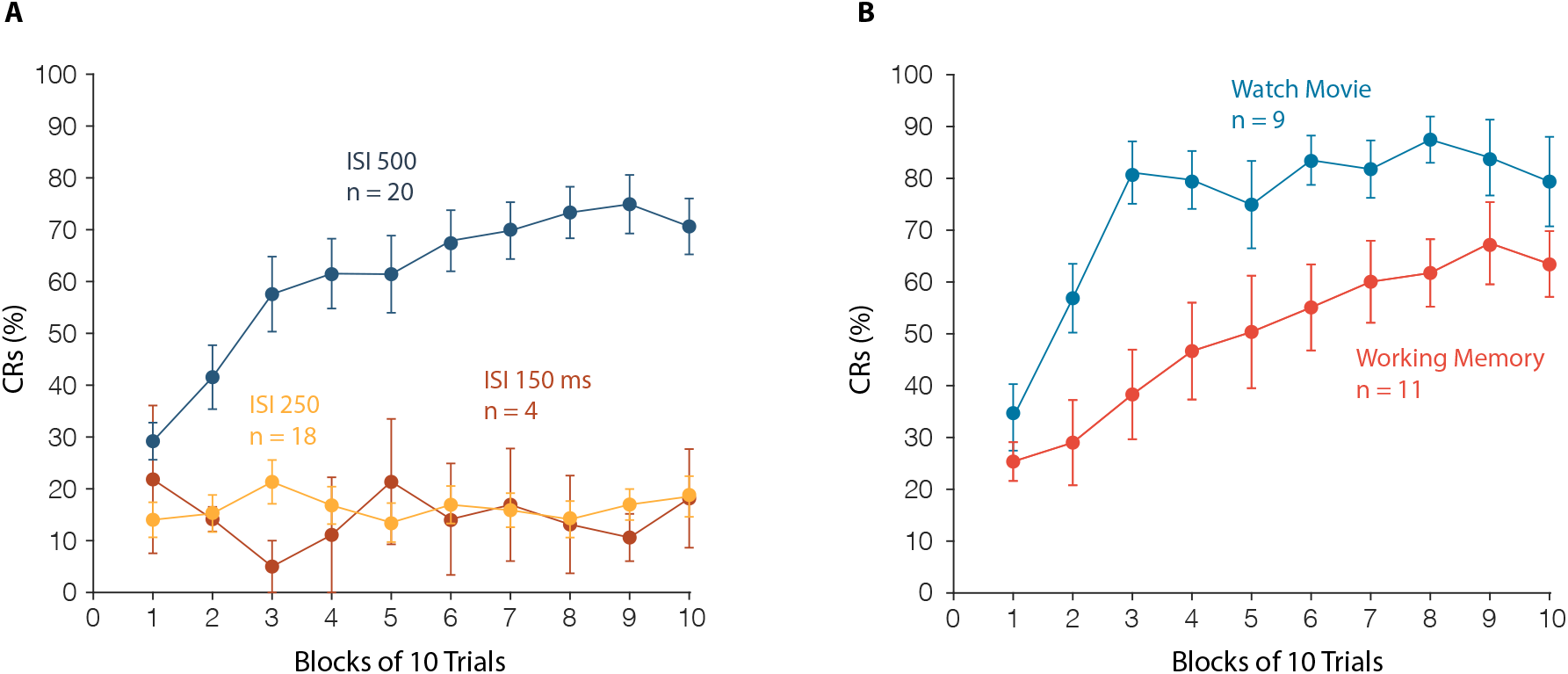
Effect of training, ISI, and working memory tasks on percentage of CRs. **A.** Displays the percentage of CRs over 10 blocks of training for subjects trained with a CS-US interval of 150 ms (yellow), 250 ms (red), and 500 ms (blue). **B.** Displays the percentage of CRs over 10 blocks of training for subjects trained with a CS-US of 500 ms who either did (red) or did not (blue) perform working memory tasks during the conditioning.

No subjects in the 500 WM group reported awareness of the CS-US contingencies, but conditioned responses were still successfully acquired. Explicit awareness was thus not necessary for successful conditioning. Whether or not the subjects solved working memory tasks during the conditioning had an effect on the CR percentage (illustrated in Figure 1B). Those who performed the tasks produced 13.68% fewer CRs than those who watched a film. This difference was statistically significant (t = 2.77, p = 0.0057**, CI = 3.99 – 23.4%). In a separate model, which included only subjects that did the working memory tasks, we tested whether a subject’s relative ranking on working memory performance predicted the percentage of CRs. The results showed that this was not the case (t = 1.3189, p = 0.18, CI = −0.9 – 5.0%). In summary the results show that: (1) training results in more CRs; (2) a longer CS-US interval results in more CRs; (3) sex does not, in this case, have an effect on CR percentage; (4) doing a working memory task reduces the percentage of CRs; and (5) how well subjects perform on the working memory tasks do not affect the CR percentage.

## Discussion

Short CS-US intervals (150 ms and 250 ms) did not induce learning. This result is markedly different from the animal literature where CS-US intervals between 150 ms and 500 ms typically produce high rates (80-100%) of conditioned responses [13,14,18–20]. This may mean that using short CS-US intervals is not a viable strategy to induce conditioned eyeblinks in humans that are comparable to the animal literature, i.e., learned blinking that primarily depends on plasticity mechanisms in the cerebellum.

Training with a 500ms ISI did induce learning. In the control group (without the working memory distraction) there was a rapid increase in rates of responding. In fact, already within the first three blocks the response rate reached >80%, and did not change much for the remaining seven blocks. This is a common observation in eyeblink conditioning with human subjects [9,10,12,21–24]. This is again markedly different from conditioning in other mammals. For example, in intact rabbits, rats and mice, conditioned responses are sometimes seen only after several days of training [3,14,18,19,25]. When responses start appearing, rates are low. Peak rates of responding and plateauing of the average learning curve usually takes additional days of training.

In the current experiments, the use of concurrent working memory tasks during training with the 500 ms CS-US interval had the effect of rendering subjects unaware of the stimulus contingencies, while still producing high rates of responding at the end of the session. Importantly, their learning curve did not plateau early, as in the control group. The gradual increase in conditioned responses extended over nine of the ten blocks of training. Also, though learning was more gradual and the rates of responding were lower from the first block and on, the response rate eventually reached the same level (>80%) as the control group by the end of the ten blocks of training.

Awareness of the stimulus conditions and voluntary blinking in response to the conditional stimulus may have been a source of increased blinking already in the early stages of learning in the 500 ms non-WM group (i.e. the control group). It may be the case that using concurrent working memory tasks during conditioning is a useful method to eliminate the problem of cerebrum dependent voluntary blinking and that it may be the case that the resulting conditioned eyeblink responses are in fact cerebellar dependent and thus more comparable to conditioned blink responses reported in the animal literature.

## Acknowledgements

PsyToolkit is developed by and belongs to Professor Gijsbert Stoet.

## References

1. McCormick DA, Thompson RF. Ceresbellum: essential involvement in the classically conditioned eyelid response. Science. 1984;223:296–9.

2. Yeo CH, Hardiman MJ, Glickstein M. Discrete lesions of the cerebellar cortex abolish the classically conditioned nictitating membrane response of the rabbit. Behavioral Brain Research. 1984;13:261–6.

3. Ten Brinke MM, Boele H-J, Spanke JK, Potters J-W, Kornysheva K, Wulff P, et al. Evolving Models of Pavlovian Conditioning: Cerebellar Cortical Dynamics in Awake Behaving Mice. Cell Rep. Elsevier; 2015;13:1977–88.

4. Jirenhed D-A, Hesslow G. Are Purkinje Cell Pauses Drivers of Classically Conditioned Blink Responses? Cerebellum. Springer-Verlag; 2016;15:526–34.

5. Zucca R, Rasmussen A, Bengtsson F. Climbing Fiber Regulation of Spontaneous Purkinje Cell Activity and Cerebellum-Dependent Blink Responses. eNeuro [Internet]. 2016;3.

6. Freeman JH, Steinmetz AB. Neural circuitry and plasticity mechanisms underlying delay eyeblink conditioning. Learn Mem. Cold Spring Harbor Laboratory Press; 2011;18:666–77.

7. Norman RJ, Villablanca JR, Brown KA, Schwafel JA, Buchwald JS. Classical eyeblink conditioning in the bilaterally hemispherectomized cat. Exp Neurol. 1974;44:363–80.

8. Weiss C, Bouwmeester H, Power JM, Disterhoft JF. Hippocampal lesions prevent trace eyeblink conditioning in the freely moving rat. Behav Brain Res. 1999;99:123–32.

9. Blaxton TA, Zeffiro TA, Gabrieli JD, Bookheimer SY, Carrillo MC, Theodore WH, et al. Functional mapping of human learning: a positron emission tomography activation study of eyeblink conditioning. Journal of Neuroscience. 1996;16:4032–40.

10. Löwgren K, Bååth R, Rasmussen A, Boele H-J, Koekkoek SKE, De Zeeuw CI, et al. Performance in eyeblink conditioning is age and sex dependent. PLoS One. 2017;12:e0177849.

11. Kjell K, Löwgren K, Rasmussen A. A Longer Interstimulus Interval Yields Better Learning in Adults and Young Adolescents. Front Behav Neurosci. 2018;12:299.

12. Kimble GA, Mann LI, Dufort RH. Classical and instrumental eyelid conditioning. J Exp Psychol. American Psychological Association; 1955;49:407–17.

13. Rasmussen A, Ijpelaar ACHG, De Zeeuw CI, Boele H-J. Caffeine has no effect on eyeblink conditioning in mice. Behav Brain Res. Elsevier B.V.; 2018;337:252–5.

14. Schneiderman N, Gormezano I. Conditioning of the nictitating membrane of the rabbit as a function of the CS-US interval. J Comp Physiol Psychol. 1964;57:188–95.

15. Rasmussen A, Jirenhed D-A. Learning and Timing of Voluntary Blink Responses Match Eyeblink Conditioning. Sci Rep. Nature Publishing Group; 2017;7:3404.

16. Krueger C, Tian L. A comparison of the general linear mixed model and repeated measures ANOVA using a dataset with multiple missing data points. Biol Res Nurs. 2004;6:151–7.

17. Dijkers MP. Chasing change: repeated-measures analysis of variance is so yesterday! Arch Phys Med Rehabil. 2013;94:597–9.

18. Kehoe EJ, Macrae M. Fundamental behavioral methods and findings in classical conditioning. In: Moore JW, editor. A neuroscientist’s guide to classical conditioning. New York: Springer-Verlag; 2002. p. 171–231.

19. Heiney SA, Wohl MP, Chettih SN, Ruffolo LI, Medina JF. Cerebellar-Dependent Expression of Motor Learning during Eyeblink Conditioning in Head-Fixed Mice. Journal of Neuroscience. 2014;34:14845–53.

20. Freeman JH, Spencer CO, Skelton RW, Stanton ME. Ontogeny of eyeblink conditioning in the rat: Effects of US intensity and interstimulus interval on delay conditioning. Psychobiology. US: Psychonomic Society; 1993;21:233–42.

21. Ebel HC, Prokasy WF. Classical eyelid conditioning as a function of sustained and shifted interstimulus intervals. J Exp Psychol. US: American Psychological Association; 1963;65:52–8.

22. Solomon PR, Pomerleau D, Bennett L, James J, Morse DL. Acquisition of the classically conditioned eyeblink response in humans over the life span. Psychol Aging. 1989;4:34–41.

23. Allen MT, Myers CE, Williams D, Servatius RJ. US alone trials presented during acquisition do not disrupt classical eyeblink conditioning: Empirical and computational findings. Behav Brain Res. Elsevier B.V.; 2018;338:101–8.

24. Wu B, Yao J, Wu G-Y, Li X, Gao W-J, Zhang R-W, et al. Absence of associative motor learning and impaired time perception in a rare case of complete cerebellar agenesis. Neuropsychologia. 2018;117:551–7.

25. Albergaria C, Silva NT, Pritchett DL, Carey MR. Locomotor activity modulates associative learning in mouse cerebellum. Nat Neurosci. Nature Publishing Group; 2018;21:725–35.

